# Gradual development of non-adjacent dependency learning during early childhood

**DOI:** 10.1101/2020.09.01.277822

**Authors:** Mariella Paul, Claudia Männel, Anne van der Kant, Jutta L. Mueller, Barbara Höhle, Isabell Wartenburger, Angela D. Friederici

## Abstract

In order to become proficient native speakers, children have to learn the morpho-syntactic relations between distant elements in a sentence, so-called non-adjacent dependencies (NADs). Previous research suggests that NAD learning in children comprises different developmental stages, where until 2 years of age children are able to learn NADs associatively under passive listening conditions, while starting around the age of 3 to 4 years children fail to learn NADs during passive listening. To test whether the transition between these developmental stages occurs gradually, we tested children’s NAD learning in a foreign language using event-related potentials (ERPs). We found ERP evidence of NAD learning across the ages of 1, 2 and 3 years. The amplitude of the ERP effect indexing NAD learning, however, decreased with age. These findings might indicate a gradual transition in children’s ability to learn NADs associatively. Cognitively, this transition might be driven by children’s increasing knowledge of their native language, hindering NAD learning in novel contexts. Neuroanatomically, maturation of the prefrontal cortex might play a crucial role, promoting top-down learning, affecting bottom-up, associative learning. In sum, our study suggests that NAD learning under passive listening conditions undergoes a gradual transition between different developmental stages during early childhood.

**Research Highlights:**

- Transition between developmental stages of non-adjacent dependency (NAD) learning
- Children between 1 and 3 years of age showed learning of NADs in a foreign language
- Brain responses revealed associative NAD learning, triggered by passive listening
- Gradual decrease of the strength of associative non-adjacent dependency learning

## 1 Introduction

In order to successfully communicate with their environment, infants must not only learn the words of their native language(s) but also the relations that define how these words combine into phrases and sentences. These relations can hold for neighboring elements in a sentence (e.g., *<u>Mary is</u> happy*) or non-neighboring elements (e.g., *The sister <u>is</u> sing<u>ing</u>*). These latter relations, so-called non-adjacent dependencies (NADs), require the learner to track dependent elements (*is* and –*ing* in these examples) across one or more intervening elements (here: *sing-*). NADs are a crucial aspect of natural languages and are present, for example, in subject-verb agreement and English tense marking. Nevertheless, NADs are relatively difficult to learn and behavioral evidence indicates that, during both natural language acquisition and artificial language learning, children learn NADs only in their second year of life, around 14-15 months of age, depending on the acquired language (Culbertson et al., 2016; Gómez & Maye, 2005; Höhle et al., 2006; Santelmann & Jusczyk, 1998).

The learning of NADs during early childhood has been shown to undergo different developmental stages, both in terms of behavioral and neurophysiological learning measures (Culbertson et al., 2016; Mueller et al., 2019; van der Kant et al., 2020). For example, Culbertson and colleagues (2016) investigated French infants’ learning of NADs in subject-verb agreement in French stimuli and observed two stages of different behavioral responses in a head-turn preference procedure. Across age, children displayed two cycles of a shift from a familiarity preference for encountered NADs to a novelty preference. The authors proposed that the processes related to the first stage, from 14 to 18 months of age, reflect initial surface-level representations based on *phonological* features of the NADs. In contrast, the processes related to the second stage, from 21 to 24 months of age, were interpreted as higher-level representations of the *morphological* features (Culbertson et al., 2016). The current paper aims to investigate the transition between these developmental stages in NAD learning more closely by looking at the development of surface-level phonological NAD learning in non-native speech stimuli throughout early childhood using event-related potentials (ERPs).

Previous studies using ERPs have provided evidence for the existence of different developmental stages of NAD learning. Friederici and colleagues (2011) tested German 4-month-old infants’ NAD learning in a passive listening familiarization-test paradigm using a non-native language (Italian) containing NADs (e.g. *La sorella <u>sta</u> cant<u>ando</u>*; *The sister <u>is</u> sing<u>ing</u>*). By means of ERPs, the authors showed that even at the early age of 4 months, infants succeeded at NAD learning. Learning was indexed by a late positive ERP effect, which was interpreted as associative learning of NADs based on phonological cues (Friederici et al., 2011). NAD learning in this miniature version of Italian and in similarly structured syllable sequences was shown to display different developmental stages during early childhood, with an initial stage up to the age of 2 to 3 years, during which young children can learn NADs through passive listening, which likely triggers associative learning (Friederici et al., 2011; Mueller et al., 2019; van der Kant et al., 2020). This is followed by a stage starting at around 3 to 4 years, during which children and adults struggle to learn NADs under passive listening (without additional cues marking the NADs), but succeed in learning under active listening conditions, that is, when a task, administered during the whole experiment or during testing only, guides their attention towards the NADs (Friederici et al., 2013; Lammertink et al., 2019; Mueller et al., 2012; Mueller et al., 2019; see also Pacton et al., 2015; Pacton & Perruchet, 2008). Notably, adults’ neural signature of NAD learning under active listening conditions differed from infants’ neural signature under passive listening: Adults under active task conditions showed an N400-like, negative ERP component and a P3, which were interpreted as indicating lexicalization of the NADs and attention-driven processing respectively (Mueller et al., 2009). This is in contrast to infants’ associative learning triggered by passive listening, which was based on phonological information processing, indexed by a late positive ERP component (Friederici et al., 2011, 2013). This difference in ERP responses between adults and infants was proposed to partly be driven by the maturation of the prefrontal cortex which supports top-down processes in contrast to the temporal cortex subserving bottom-up associative processes (Skeide & Friederici, 2016). This proposal is in line with the results of a study that tested adults’ NAD learning while top-down processing was suppressed (by inhibiting the prefrontal cortex with transcranial direct current stimulation; Friederici et al., 2013). Under these conditions, adults showed a late positive ERP component in response to NAD violations, comparable to the ERP response found in infants using the same paradigm. This ERP response was interpreted to indicate associative NAD learning, comparable to infants’ learning (Friederici et al., 2013). Thus, ERP research on NAD learning further corroborates the observation of different developmental stages, as the neural signature of NAD learning seems to change with age, and is dependent on the presence or absence of a task. However, it is less clear exactly when or how the transition between these developmental stages occurs, and whether the transition occurs abruptly or more gradually across preschool age.

Investigating the nature of the transition between the different developmental stages of NAD learning can improve our understanding of children’s acquisition of grammatical rules and language learning in general. A rich body of literature has demonstrated that the ability for language learning, and in particular learning grammatical rules, has its peak in infancy and decreases over development (e.g., Hartshorne et al., 2018; Johnson & Newport, 1989; Senghas et al., 2004; Singleton, 2005). It has been suggested that this time course is driven by a sensitive period for language learning under passive listening conditions influenced by brain plasticity (e.g. Knudsen, 2004; Kuhl, 2004, 2010; see also Skeide & Friederici, 2016). Crucially, first language learning occurs under passive listening conditions. That is, infants learn NADs in their first language by simply listening to their environment, without being taught explicit rules (see Perruchet & Pacton, 2006). If NAD learning under passive listening conditions indeed undergoes a sensitive period, defined by a gradual closing of the period (Knudsen, 2004), we would expect to see a gradual transition between the developmental stages of NAD learning. Based on the studies reviewed above, we would expect to observe surface-based learning of phonological associations during this potential sensitive period.

In the present study, we aimed to confirm the presence of different developmental stages of NAD learning found in previous studies (Culbertson et al., 2016; Mueller et al., 2019; van der Kant et al., 2020) and, for the first time, systematically investigate the nature of the transition between the different developmental stages of NAD learning under passive listening conditions. In particular, we aimed to investigate whether this transition occurs in a gradual manner, which may point towards a sensitive period of NAD learning under passive listening conditions. To investigate this, we used the same familiarization-test paradigm with Italian sentences as previous ERP studies that provided evidence for different developmental stages of NAD learning (Friederici et al., 2011, 2013; Mueller et al., 2009; van der Kant et al., 2020). Using a set of non-native sentences allows for combining the advantages of artificial grammars, by controlling for children’s exposure to the language, and those of natural language, by being more naturalistic than artificial grammars. Based on previous findings with the same paradigm and artificial grammar paradigms, we propose that a transition between different developmental stages of NAD learning takes place between 2 and 4 years of age (Mueller et al., 2019; van der Kant et al., 2020). To investigate whether this transition occurs in a gradual manner, that is, whether we can observe a linear decline in NAD learning under passive listening conditions during early childhood, we exposed 1-to 3-year-old children to Italian sentences under passive listening conditions in a familiarization-test paradigm while recording ERPs. ERPs offer the advantage that they can be measured throughout the age range tested here, because they do not require children to give explicit responses or to exhibit any kind of behavior (as in the head-turn preference procedure). ERPs also allow us to make inferences based on both the strength (amplitude) and the timing (latency) of children’s ERP responses. Based on previous infant and adult ERP studies using the same familiarization-test paradigm (Friederici et al., 2011, 2013), we expected passive listening to trigger associative learning. In these previous studies, detection of NAD violations was indexed by a late positive ERP effect in response to incorrect sentences (containing NAD violations) compared to correct sentences (containing familiarized NADs) during test phases. If the transition between the different stages of NAD learning indeed occurs in a gradual manner, we would expect to see a linear decrease in the amplitude of this ERP effect of NAD learning between the age of 1 and 3 years. If the transition instead occurs more abruptly, we would not expect to see a linear decrease in ERP amplitude between the ages of 1 and 3 years; in this case we would expect the amplitude of the ERP effect to be either unchanged across the ages tested here or to show evidence for learning in the younger, but not in the older children. More specifically, we hypothesize a significant difference in ERP responses to NAD violations compared to familiarized NADs under passive listening conditions for 1-year-old children, who have been shown to successfully learn NADs in an associative manner (Culbertson et al., 2016). For 2-year-olds, we expect successful NAD learning under passive listening conditions (Mueller et al., 2019; van der Kant et al., 2020), although the strength of associative learning under passive listening conditions may have started to decrease (Culbertson et al., 2016). Finally, for 3-year-old children, we expect a further decline of the strength of NAD learning under passive listening conditions, based on a previous ERP study that found that only a subgroup of 4-year-old children learned NADs under passive listening (Mueller et al., 2019, but see van der Kant et al., 2020 for no evidence for NAD learning under passive listening conditions at 3 years of age using functional near-infrared spectroscopy (fNIRS).

In previous ERP studies on NAD learning, the detection of NAD violations was indexed by different ERP responses, depending on the specifics of the experimental design and stimuli as well as the studied participant sample. In particular, in the studies using the same Italian sentence material as we use here, the ERP polarity differed depending on participants’ age and whether participants were tested under passive listening conditions or performed a violation detection task (Friederici et al., 2011, 2013; Mueller et al., 2009). Moreover, ERP polarity was shown to change with the discrimination difficulty of the chosen stimuli (Schaadt & Männel, 2019), infants’ sex (Mueller et al., 2012), and whether school-age children were tested immediately after NAD learning or following a retention period involving sleep (Schaadt et al., 2020). Similarly, the polarity of infants’ ERP effects was found to be associated with later language outcomes in studies testing lexical segmentation and phoneme discrimination (Kooijman et al., 2013; Schaadt et al., 2015). Based on the polarity differences in these studies, we do not make a priori predictions about the polarity of the measured ERP effects, but investigate whether it plays a role in the developmental stages of NAD learning and the transition between them. We included ERP polarity in an exploratory analysis and investigated whether the ERP effect’s polarity was associated with children’s age, sex or behaviorally tested language development (comprehension and production), as it was in previous ERP studies (Friederici et al., 2011, 2013; Kooijman et al., 2013; Mueller et al., 2012; Schaadt et al., 2015).

## 2 Material and Methods

### 2.1 Participants

115 healthy children growing up in monolingual German families participated in this study. Of these, 40 were 1 year old (mean age: 12.80 months, SD: 0.54; 20 girls), 40 were 2 years old (mean age: 25.08 months, SD: 0.88; 16 girls) and 35 were 3 years old (mean age: 37.10, SD: 0.60; 18 girls). Children were orally informed about the experimental procedure and caregivers were informed both in written and oral form. Caregivers gave written informed consent for their children’s participation in the study. Ethical approval was obtained from the Medical Faculty of the University of Leipzig. Forty-nine additional children had to be excluded from data analysis, either due to non-compliance during the experimental procedure (22 children) or because they contributed less than 10 artefact-free EEG trials per condition across all test phases and/or no trials in at least one test phase (27 children). For additional information on EEG trial numbers see Table 1.

**Table 1.** Overview of participants and trials. “Additional participants” refers to children that were tested, but excluded based on either non-compliance or insufficient number of artefact-free EEG trials. Correct and incorrect trials refer to the average number of included trials in the correct and incorrect condition during test phases. The difference in trial numbers between the correct and incorrect conditions was not significant for any age group (all p>0.1).

|  | 1 year | 2 years | 3 years |
| --- | --- | --- | --- |
| Participants (N) | 40 | 40 | 35 |
| Mean age (months) | 12.80 (SD: 0.54) | 25.08 (SD: 0.88) | 37.10 (SD: 0.60) |
| Sex | 20 girls | 16 girls | 18 girls |
| Additional participants (N) | 28 | 15 | 6 |
| Correct Trials (N) | 21.90 (SD: 4.24) | 24.25 (SD: 4.47) | 26.77 (SD: 4.15) |
| Incorrect Trials (N) | 22.53 (SD: 3.85) | 23.93 (SD: 4.70) | 27.26 (SD: 3.59) |

### 2.2 Stimuli

#### 2.2.1 EEG experiment

The stimuli for the EEG experiment were adapted from Mueller and colleagues (2009). Participants listened to Italian sentences (see Figure 1) spoken by a female native speaker of Italian. The sentences consisted of a determiner phrase (*La sorella* (*The sister*) or *Il fratello* (*The brother*)), followed by a verb phrase consisting of an auxiliary (*sta* (*is*) or a modal verb *puo* (*can*)), a verb stem (32 different verbs, e.g., *cant-* (*sing-*)), and a suffix (*-ando* (*-ing*) or *-are* (*-*∅)). In grammatical sentences, *sta* (*is*) is followed by a verb stem and *-ando* (*-ing*), while *puo* (*can*) is followed by a verb stem and *-are* (*-*∅). In ungrammatical sentences, the verb suffixes did not match the auxiliary, namely *sta* (*is*) was followed by a verb stem and *-are* (*-*∅), while *puo* (*can*) was followed by a verb stem and *-ando* (*-ing*). Crucially, to avoid introducing acoustic cues regarding the grammaticality of the sentence as well as potential acoustic differences between the two conditions, the speaker produced only grammatical sentences and both correct and incorrect sentences underwent a cross-splicing procedure (Adobe Audition), in which the verb was exchanged with the verb from a different sentence (see Friederici et al., 2011). Sentences had a mean length of 2.43 s (SD: 0.12). Because we counterbalanced whether participants were familiarized with grammatical or ungrammatical Italian sentences, we will refer to the NADs presented in the familiarization phases as correct (i.e., regardless of grammaticality in Italian).

**Figure 1.**
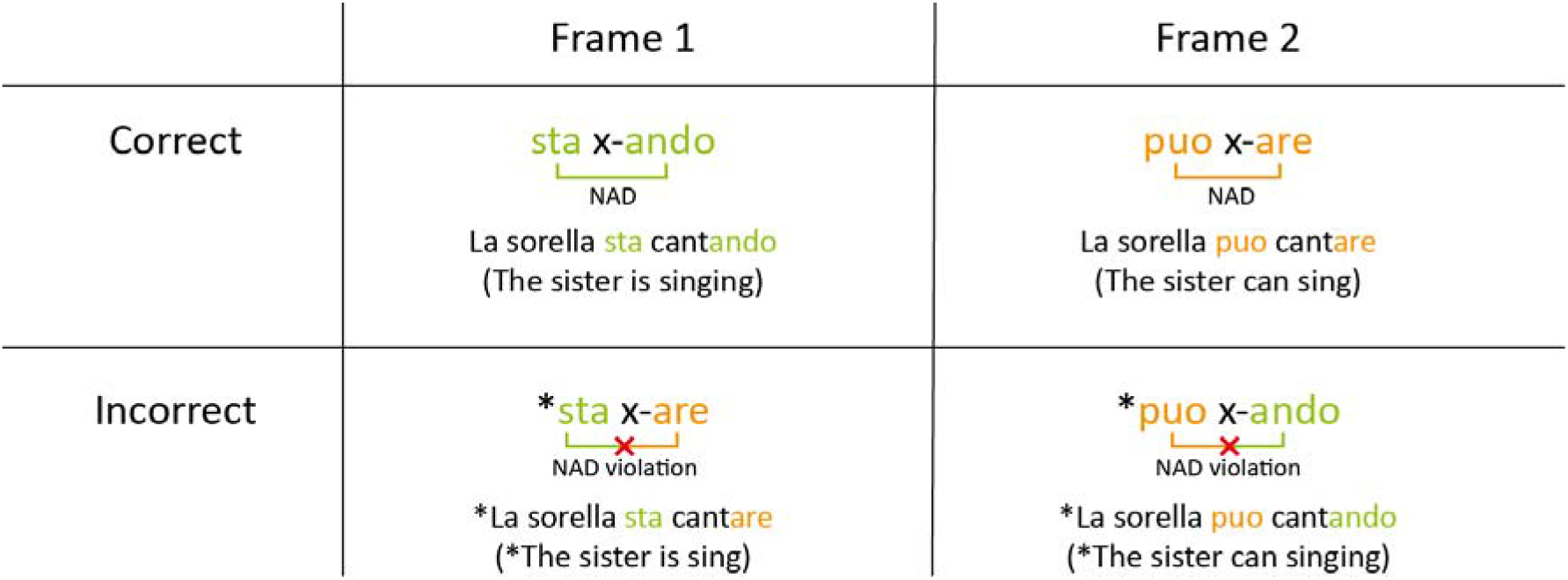
Visualization of the stimuli for the EEG experiment with examples, adapted from Friederici and colleagues (2011). Ungrammatical sentences and frames are marked with an asterisk. Unicolored brackets visualize non-adjacent dependencies (NADs). Bicolored brackets and red crosses indicate NAD violations.

### 2.2.2 Tests of language development

For all three age groups, we behaviorally assessed language abilities via standardized tests. For 1-year-old children, we used the German version of the *Bayley Scales of Infant and Toddler Development Screening Tes*t (Bayley, 2015). For the language comprehension measure, we used the subscale receptive language of the Screening Test and for the language production measure, we used the subscale productive language. Both of these subscales assess children’s vocabulary in a playful interactive manner. For 2- and 3-year-old children, we used the *Sprachentwicklungstest für zweijährige Kinder* (SETK-2; Grimm et al., 2016) and *Sprachentwicklungstest für drei-bis fünfjährige Kinder* (SETK 3-5; Grimm, 2015), respectively. For 2-year-old children’s language scores, we used the averaged word and sentence comprehension and production subscales. For 3-year-old children’s language scores, we used the sentence comprehension and production measures. Because the Bayley Screening Test only offers raw scores, we used z-transformed raw scores of all tests to allow for comparisons across age groups.

### 2.3 Procedure

During the EEG experiment, participants were seated on their caregiver’s lap in a soundproof booth. The experiment consisted of four familiarization phases alternating with four test phases. Familiarization phases (3.3 minutes each) consisted of 64 correct sentences each (overall 256 sentences). Each familiarization phase was followed by a test phase (1.3 minutes each), consisting of 8 correct and 8 incorrect sentences each (overall 32 sentences per condition; see Figure 2). Note that the sentences that participants heard in the test phases were not repeated in any of the familiarization phases. Furthermore, verb stems were divided into two sets, such that they were not presented in the familiarization phase immediately preceding a given test phase, but only occurred in earlier or later familiarization phases. The inter-stimulus-intervals (ISI) were 580 ms in the familiarization phases and 1380 ms in the test phases. Between familiarization and test phases, there was a pause of 2780 ms. Overall, the EEG experiment took approximately 20 min, during which participants watched a silent children’s movie (*Peppa Pig, Bummi*, or *Alles Trick 9*) in order to increase compliance. The EEG experiment and the standardized language tests were either administered during the same session or in two separate sessions (mean time between sessions: 13.35 days, SD: 14.78).

**Figure 2.**
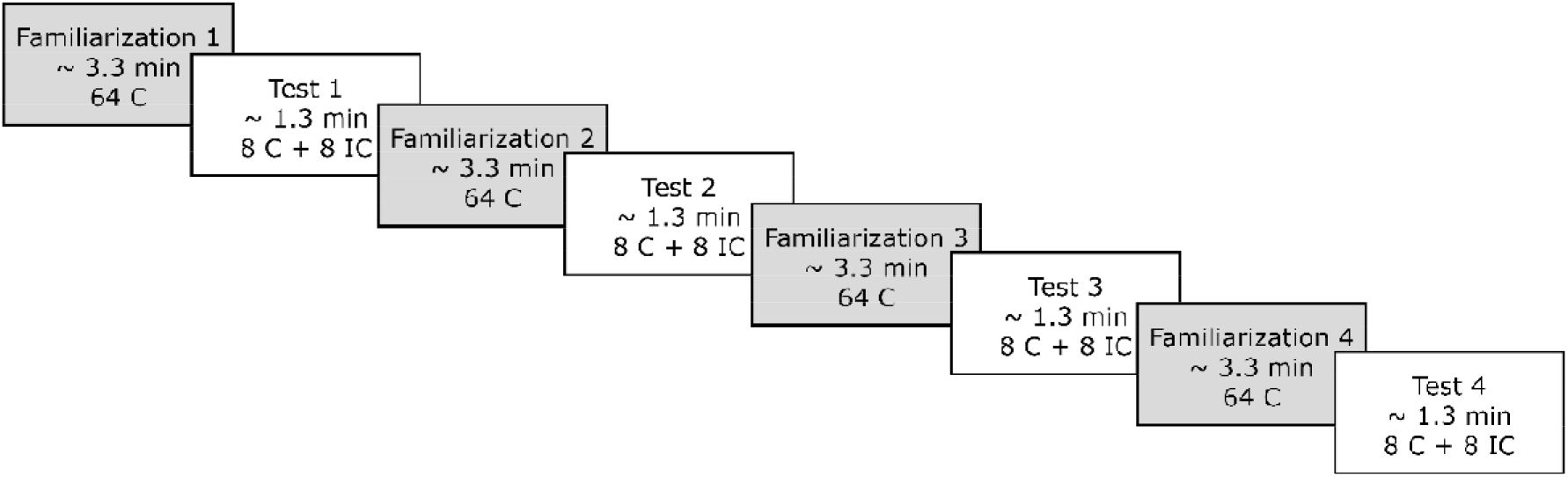
Experimental procedure: alternating familiarization and test phases. Participants listened to four familiarization phases (64 correct trials each) alternated with four test phases (eight correct and eight incorrect trials each). C: correct, IC: incorrect. Figure adapted from Friederici, Mueller, and Oberecker (2011).

### 2.4 Data recording

EEG data were recorded from 24 Ag/AgCl electrodes (Fp1, F7, F3, Fz, F4, F8, FC3, FC4, T7, C3, Cz, C4, T8, CP5, CP6, P7, P3, Pz, P4, P8, O1, O2, M1, and M2) placed according to the International 10-20 System of Electrode Placement and secured in an elastic electrode cap (Easycap GmbH, Herrsching, Germany). Cz served as an online reference during recording. Electrooculograms (EOG) were recorded from 4 additional electrodes, placed supraorbitally (Fp2) and infraorbitally (V-) to the right eye to capture vertical eye movements, as well as laterally to the left (F9) and right eye (F10), for horizontal eye-movements. The signal was digitized with a sampling rate of 500 Hz.

The EEG data were analyzed using the Fieldtrip toolbox (Oostenveld et al., 2011) implemented in Matlab (MATLAB, 2017). Scripts for both, preprocessing and analysis can be found on the Open Science Framework (https://osf.io/43t9q). The signal was re-referenced offline to the linked mastoids (the algebraic average of M1 and M2) and down-sampled to 250 Hz. We applied a kaiser-windowed finite-impulse response high-pass filter with half-amplitude cutoff (−6 dB) of 0.3 Hz and a transition width of 0.3 Hz. We also applied a kaiser-windowed finite-impulse response low-pass filter with a half-amplitude cutoff (−6dB) of 30 Hz and a transition width of 5 Hz. Data were segmented and time-locked to the onset of the suffix, with a 600 ms pre-stimulus period (to include the onset of the verb stem) and 1300 ms post-suffix-onset period. Artefact rejection was performed semi-automatically. Segments of the signal exceeding a z-value of 7 were highlighted automatically and screened manually to reject muscle and coarse-movement artefacts. To correct ocular artefacts, we used an independent component analysis (ICA) (“runica”, implemented in the FieldTrip toolbox; Oostenveld et al., 2011), decomposed the data from all channels into 26 ICA components, and rejected components corresponding to blinks and saccades. Afterward, we shortened the baseline period to 400 ms pre-suffix onset for plotting and averaging.

### 2.5 Statistical analysis

For the main statistical analysis, we used linear models (LMs; see Frömer et al., 2018 for use of LMs and linear mixed models with EEG data), as implemented in the lme4 package (Bates et al., 2015) in R (R Core Team, 2017). LMs offer a reliable way to analyze all three age groups in one statistical model. The time window of interest was defined as 600-1000 ms relative to the onset of the suffix (-are, -ando) and the spatial region of interest included the electrode positions F3, Fz, F4, C3, Cz, C4, P3, Pz, P4, based on a previous ERP study with the same paradigm in infants (Friederici et al., 2011). Thus, we computed mean amplitudes across the defined time window and region of interest for each subject.

#### 2.5.1 Predictors of ERP polarity

Fifty-seven percent of the participants (N=66) showed a positive-going ERP effect (correct vs. incorrect) in the time window and region of interest, while the other 43% (N=49) showed a negative-going ERP effect. We here used a logistic regression to investigate whether children’s ERP polarity (positive vs. negative) could be significantly predicted by their age (1 year, 2 years, 3 years; using linear contrast coding; see Schad et al. (2020) for further information on contrast coding), sex (male, female; using sum contrast coding), or language comprehension or language production abilities (z-transformed raw scores of the language tests). This analysis was performed in R (R Core Team, 2017).

#### 2.5.2 Age effects of NAD learning

Using the time window and region of interest defined above, we set up an LM in R (R Core Team, 2017) to test for the effect of age on children’s ERP effect (correct vs. incorrect), as an indicator for NAD learning. As the dependent variable, we used the amplitude of the absolute (based on the results of the linear regression, see section 3.1) ERP difference wave (incorrect – correct; see Figure 3) averaged over the time window and region of interest, further averaged over trials in order to increase the signal-to-noise ratio for the LM, resulting in one ERP amplitude value per participant. When considering the ERP amplitude values entered into the LM, there was still considerable variability (mean: 0.059; SD = 6.73; min = −31.06, max = 15.69). Therefore, we excluded outliers, defined as 2.5 times the median absolute cutoff (Leys, Ley, Klein, Bernard, & Licata, 2013). This procedure resulted in the exclusion of the datasets of 6 additional children. The results of the linear model were the same regardless of outlier exclusion. The results reported in the main text do not include outliers; for the same analyses including outliers, see supplementary materials S1. We entered *age* (1 year, 2 years, 3 years; using linear contrast coding) as a fixed effect (independent variable) into the model. We further added weights to the model, accounting for the number of trials of each average.

**Figure 3.**
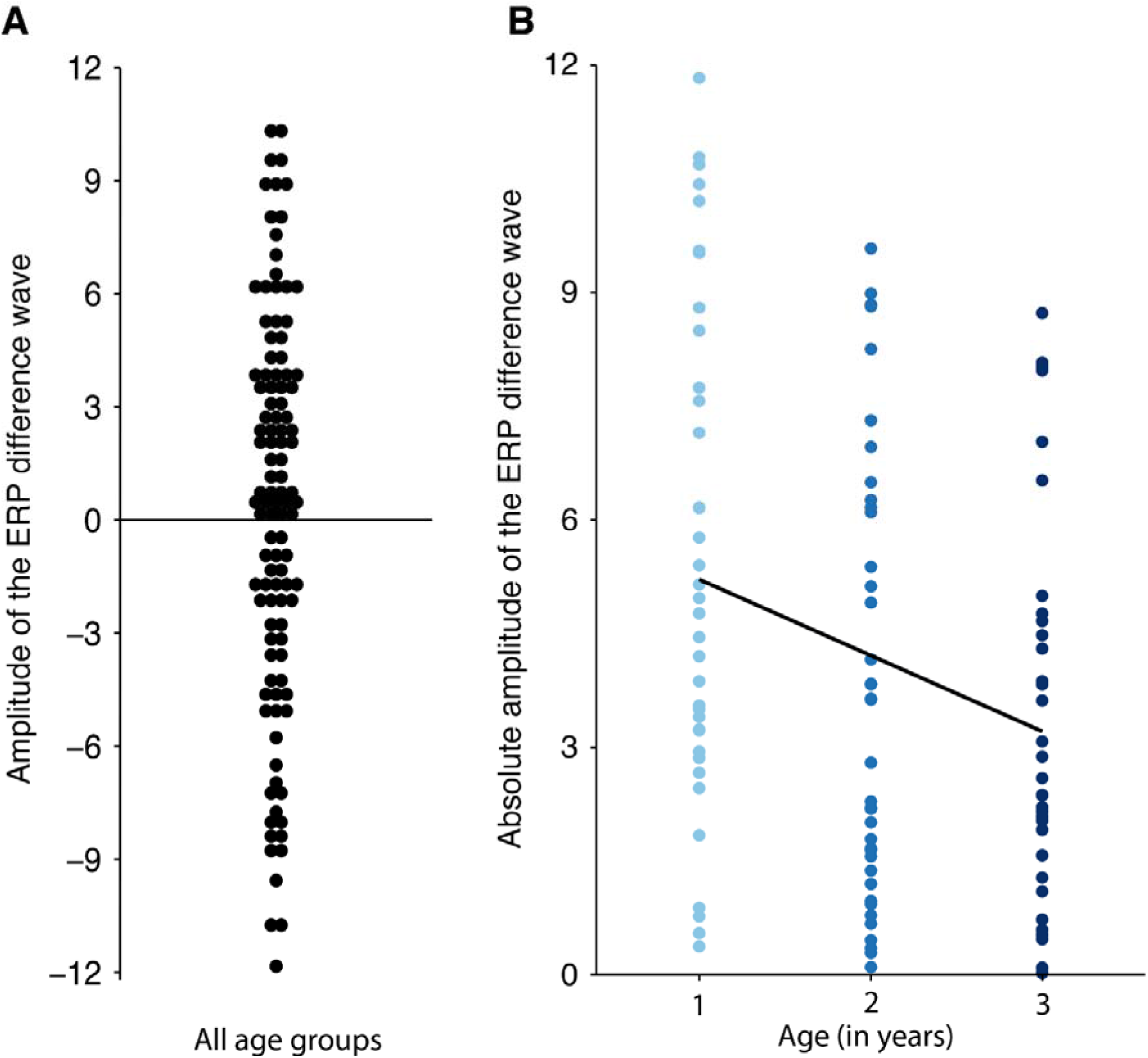
A. Amplitude of the ERP difference wave (incorrect - correct) across all age groups. B. Absolute amplitude of the ERP difference wave (incorrect - correct) plotted by age. Absolute ERP amplitude is significantly predicted by age.

#### 2.5.3 Exploratory analysis of ERP latency

To explore whether the onset and duration of the ERP effect differed between age groups we did an additional analysis on the latency of the ERP effect. This analysis was conducted in SPSS v.26 (*IBM SPSS Statistics*, 2019) and Matlab (MATLAB, 2017). We conducted a repeated-measures ANOVA with 2 factors: *age* (1, 2, and 3 years) and *time* (13 100-ms time windows from 0-1300 ms).

## 3 Results

### 3.1 Predictors of ERP polarity

The logistic regression analysis revealed that none of the tested variables (age, sex, language comprehension, and language production) significantly predicted whether children showed a positive or negative ERP effect polarity (all p>0.05; Table 2). Therefore, we used the absolute amplitude as the dependent measure in the LM.

**Table 2.** Summary of the logistic regression to predict children’s ERP effect polarity

| *Predictors* | *Odds Ratios* | *CI* | *p* |
| --- | --- | --- | --- |
| (Intercept) | 0.71 | 0.19 – 2.71 | 0.619 |
| age | 1.03 | 0.52 – 2.05 | 0.922 |
| sex | 1.44 | 0.63 – 3.38 | 0.390 |
| lang. comp. | 1.16 | 0.71 – 1.92 | 0.565 |
| lang. prod. | 1.09 | 0.68 – 1.77 | 0.713 |
| Observations | 99 |  |  |
| R<sup>2</sup> Tjur | 0.014 |  |  |

Note: lang. comp. – language comprehension. lang. prod. – language production

### 3.2 Age effects of NAD learning

A likelihood-ratio test revealed that the LM including the fixed effect *age* explained significantly more variance than a restricted model with the factor *age* omitted (F = 6.11 p = 0.015). There was a significant main effect of *age* (β = −1.20, p = 0.013; Table 3), indicating that the absolute amplitude of the ERP effect significantly decreased with increasing age (Figure 3b; Figure 4).

**Table 3.** Summary of the LM predicting children’s ERP amplitudes. Statistically significant p-values are highlighted in bold.

| *Predictors* | *Estimates* | *CI* | *p* |
| --- | --- | --- | --- |
| (Intercept) | 4.10 | 3.55 – 4.65 | **<0.001** |
| age | -1.20 | -2.17 – -0.24 | **0.015** |
| Observations | 109 |  |  |
| R<sup>2</sup> / R<sup>2</sup> adjusted | 0.054 / 0.045 |  |  |

Note: inc – incorrect condition, cor – correct condition

**Figure 4.**
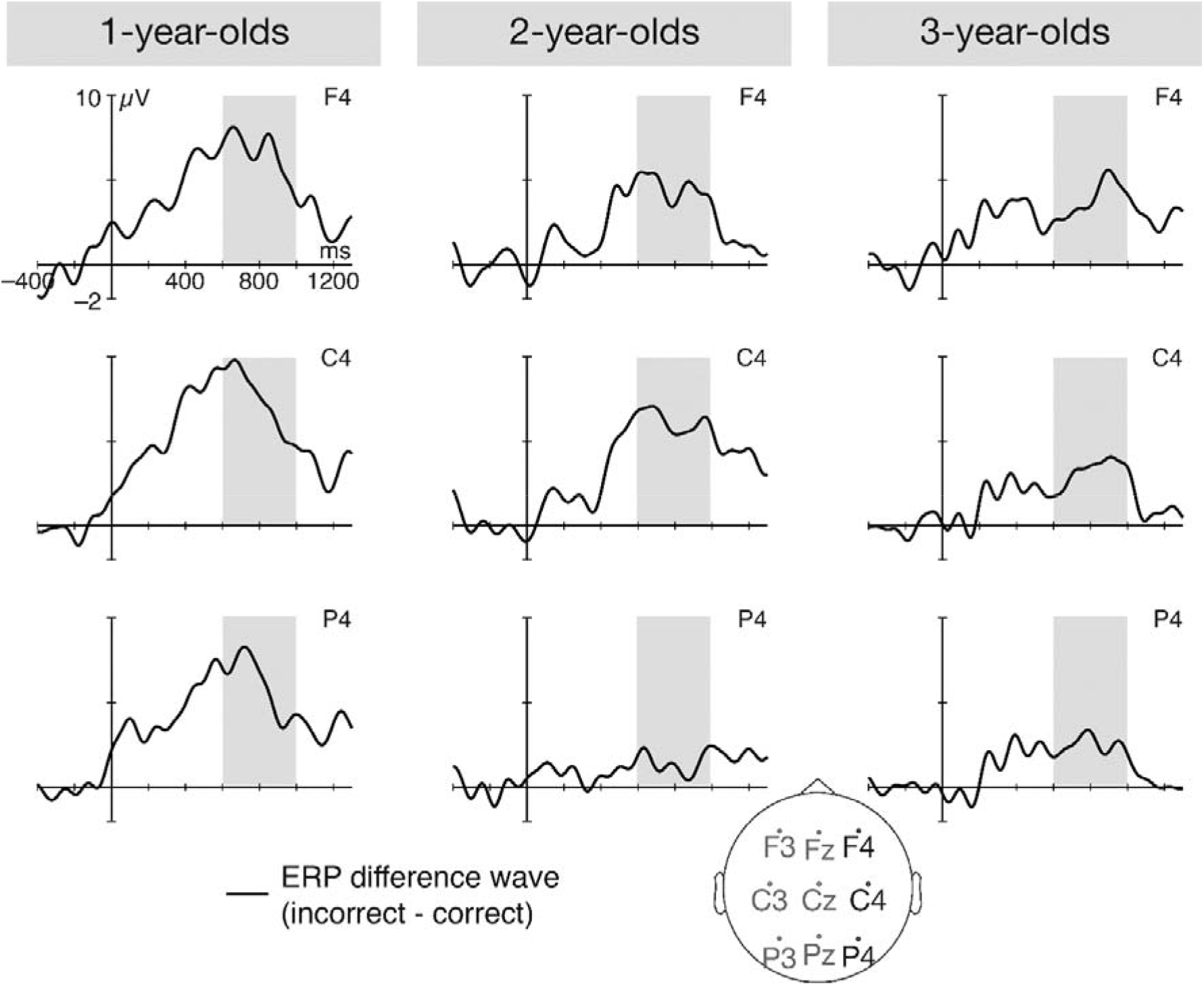
Grand-average ERP difference waves (incorrect - correct) for the three age groups. To account for the absolute amplitudes used in the analyses, the ERP difference wave for the children with negative polarity is flipped for visualization purposes. Grey bars highlight the time window from which the absolute amplitude was extracted.

We followed up on this significant effect of age with separate LMs for each age group. These LMs are equivalent to one-sample t-tests, but include weights for the number of trials that constituted each average. These one-sample t-tests revealed that for all three age groups, the absolute amplitude of the difference wave was significantly different from 0 (1 year: β = 5.23, p < 0.001; 2 years: β = 3.62, p < 0.001; β = 3.48, p < 0.001).

### 3.3 Exploratory analysis of ERP latency

The repeated-measures ANOVA with factors age (1, 2, and 3 years) and time (13 time windows of 100 ms each, ranging from 0-1300 ms) revealed a significant main effect of age (F= 4.85, p = 0.01), a significant main effect of time (F = 8.68, p < 0.001), and a marginally significant age*time interaction (F = 1.46, p = 0.063). We followed up on this interaction with individual t-tests per age group (Bonferroni-corrected for multiple testing within each age group). These t-tests revealed that for the 1-year-old children, the latency of the ERP effect was 200-1100 ms, for the 2-year-old children, it was 500-1100 ms, and for the 3-year-old children, it was 600-1000 ms. For a table of all t-values and p-values of the individual t-tests, please see Supplementary Table S2. These results indicate that increasing age is associated with a later onset as well as a shorter duration of the ERP effect.

## 4 Discussion

Previous studies have demonstrated that NAD learning undergoes different developmental stages during early childhood (Culbertson et al., 2016; Mueller et al., 2019; van der Kant et al., 2020). The aim of the present study was to investigate whether the transition between these stages occurs in a gradual or more abrupt manner. To this end, we exposed 1- to 3-year-old children to Italian sentences containing NADs and measured children’s ERP responses to incorrect sentences (containing NAD violations) compared to correct sentences (containing familiarized NADs). Independent of the tested age, children’s ERP responses <u>suggested</u> that they were able to distinguish correct from incorrect sentences, and had thus learned the NADs, indexed by an ERP component between 600 and 1000 ms (relative to suffix onset) with a positive polarity in 57% of all children and a negative polarity in 43%. Previous studies found an association of children’s ERP polarity and later language outcomes (Kooijman et al., 2013; Schaadt et al., 2015) as well as infants’ sex (Mueller et al., 2012). In addition, when the current paradigm was used in previous studies, ERP polarity had been related to participants’ age and/or the presence or absence of a task (Friederici et al., 2011, 2013; Mueller et al., 2009). Yet, neither children’s age, sex, nor their behaviorally tested language abilities did predict a given child’s ERP polarity in our study. Importantly, regardless of ERP polarity, the amplitude of the ERP effect of NAD learning decreased linearly with age, which we suggest indicates a gradual decrease in strength of NAD learning. In previous studies, ERP amplitude has been shown to be indicative of both strength of learning (Boll-Avetisyan et al., 2018) and tone and phoneme discrimination abilities (Garcia-Sierra et al., 2011; Kujala et al., 2001). A previous infant ERP study using the same paradigm as the current study found that 4-month-old infants’ NADs learning was indicated by a late positive ERP component (640-1040 ms relative to suffix onset; Friederici et al., 2011). This ERP component was interpreted to reflect associative learning, that is, infants learned associations between surface-level phonological features of the dependent elements (Friederici et al., 2011). In line with Friederici and colleagues’ (2011) <u>interpretation</u>, we propose that children in our study learned the NADs in an associative manner, via surface-level phonological features, based on the passive listening design and the findings that at least 1-year-old children are not yet able to learn the grammatical rules underlying NADs (Culbertson et al., 2016; Legendre et al., 2010). Our findings would then imply that at 1 to 3 years of age, children are in principle capable of learning NADs associatively from passive listening, but that this ability gradually decreases with increasing age. This interpretation is supported by an exploratory analysis of the latency of the ERP effect, which showed that the ERP effect starts earlier and lasts longer in younger children, indicating the effect is more robust in younger children than in older children. In line with the proposal of an age-related decrease, German adults have been shown to struggle to learn NADs from Italian sentences through passive listening (Friederici et al., 2013), but to successfully learn the NADs under active conditions, that is, in the presence of a task (Mueller et al., 2009; see also Pacton et al., 2015; Pacton & Perruchet, 2008). Together, these findings imply that during development, there is an initial stage of NAD learning, during which young children are able to learn NADs associatively, and a later stage, during which older children and adults need additional cues (e.g. Gómez, 2002; Grama & Wijnen, 2018; Newport & Aslin, 2004; for a review, see Wilson et al., 2018) or a task (Mueller et al., 2012; Pacton et al., 2015; Pacton & Perruchet, 2008) to guide their attention to successfully learn NADs. Our current findings contribute to this notion and, moreover, provide evidence that the transition between these developmental stages of NAD learning occurs in the form of a gradual decrease of associative learning during early childhood.

These findings raise the question of which processes underlie the developmental stages of NAD learning, including their transition. A behavioral study by Culbertson and colleagues (2016) proposed that an initial developmental stage of NAD learning (around 1 to 1.5 years of age) is characterized by associative learning of phonological features, while a later stage (around 2 years of age) is characterized by higher-level morphological learning. Our findings suggest that 2-year-old and even 3-year-old children are still able to learn NADs associatively from passive listening, but also indicate that children show smaller and less robust effects for associative learning with increasing age. Considering Culbertson and colleagues’ (2016) behavioral findings using the headturn-preference procedure, it is possible that the ability to learn not only the surface-level phonological features, but also the higher-level morphological features of the NADs is slowly developing between 1 and 3 years, but this learning strategy was not triggered by our passive listening task. Similarly, it is possible that different measures, such as the head-turn preference procedure compared to ERPs and different NAD learning paradigms, tap into different learning processes. This difference in measures might also explain the differences between our study and a recent study using a similar paradigm with fNIRS, in which we found NAD learning from the same Italian sentences in 2-year-old, but not 3-year-old children (van der Kant et al., 2020). While fNIRS informs us about the brain areas underlying NAD learning during early childhood, EEG may be more sensitive to detect children’s decreased responses to NAD violations at 3 years of age. Our results of a gradual decrease of associative NAD learning are therefore not necessarily at odds with previous studies reporting different developmental stages of NAD learning even before 3 years of age (Culbertson et al., 2016; van der Kant et al., 2020), but electrophysiological measures might be more sensitive to the associative processes triggered by our passive listening design.

In the following, we discuss two potential explanations for the observed gradual decrease of associative NAD learning during early childhood in the present study: (1) entrenchment of children’s knowledge of their native language, and (2) maturational brain changes during early childhood. Regarding the former, children’s early established (or entrenched) learning may influence expectations during later stages of learning (see Thiessen et al., 2016). These expectations facilitate subsequent learning of similar items, but hinder learning of new, dissimilar items. In line with this idea, entrenchment has been shown to occur and hinder learning of new items in infants’ use of lexical stress cues in word segmentation (Jusczyk et al., 1999) and in learning to read (Zevin & Seidenberg, 2002, 2004). Similarly, it is plausible that through the course of early childhood, children’s knowledge of the NADs in their native language becomes entrenched. Indeed, evidence from natural language studies show that native-language NAD learning slowly develops between 1 and 3 years of age. For example, French-learning infants can detect NAD violations in their native language starting around 14 months to 18 months, depending on the exact NADs tested (Culbertson et al., 2016; Nazzi et al., 2011; van Heugten & Shi, 2010). English-learning infants are able to detect NAD violations in ‘*is -ing*’ constructions in their native language at 18 months, but not 15 months, and only when the NADs have no more than 3 intervening syllables (Santelmann & Jusczyk, 1998; see also Höhle et al., 2006 for evidence from German-learning infants). However, detecting violations does not necessarily mean that children learn the higher-level morphological rule and comprehend the meaning of the NADs. These abilities seem to develop later, between 21 and 30 months (Culbertson et al., 2016; Legendre et al., 2010). It is <u>conceivable</u> that this increasing knowledge of children’s native language NADs makes learning NADs in a foreign language (or an artificial language, such as our miniature version of Italian) more difficult with increasing age. Indeed, infants’ NAD learning in an artificial language has been linked to processing NADs in their native language (Lany & Shoaib, 2019). This effect of entrenchment on learning novel NADs would explain why children’s ability to learn NADs associatively decreased with age in our study. Taken together, children’s knowledge of the NADs of their native language builds up over the first three years of life, possibly making learning of NADs in another language (such as our miniature version of Italian) more difficult for older children.

The second explanation of the gradual decrease in associative NAD learning refers to the maturation of the developing brain (Ramscar & Gitcho, 2007). Associative NAD learning has been proposed to demand an interplay between posterior temporal brain areas and the premotor cortex (Friederici, 2012; Gervain et al., 2008, 2011; see Skeide & Friederici, 2016). These regions are involved in language comprehension and production more generally (Bruderer et al., 2015; Möttönen & Watkins, 2009; Rodd et al., 2015) and functionally connected through ventral and dorsal fiber pathways (Dubois et al., 2016; Perani et al., 2011). The ventral pathway is already well myelinated at birth (Perani et al., 2011) and available to infants for learning NADs at a very young age, likely providing the neurobiological basis of infants’ ability to learn NADs associatively through surface-level phonological features (see Friederici et al., 2011; Skeide & Friederici, 2016). In contrast, the development of higher-level learning of morpho-syntactic NADs is likely linked to the maturation of the prefrontal cortex and the arcuate fasciculus as the dorsal pathway, connecting the posterior temporal cortex and the pars opercularis (part of the inferior frontal gyrus), which has been shown to be specifically involved in syntactic processing in adults (Rodd et al., 2015; Vigneau et al., 2006). The arcuate fasciculus, unlike the dorsal pathway which connects to the premotor cortex, shows continuous maturation until early adulthood, and has been linked to the development of syntactic processing (Skeide et al., 2016). Specifically, in young children, syntactic information triggers activation in the left temporal cortex, but not the left inferior frontal cortex; at 3 years of age, at the latest, both regions are activated during syntactic processing (Dehaene-Lambertz et al., 2002, 2006; Skeide et al., 2016). In summary of these results, Skeide & Friederici (2016) proposed a transition from associative, bottom-up learning mainly based on the temporal cortex to higher-level, top-down learning around the age of 3 years involving the left inferior frontal cortex. Further studies need to evaluate whether the *decrease* of associative learning of phonological features is accompanied by a <u>comparable</u> gradual *increase* of higher-level morphological NAD learning during the same developmental period. The interplay between a decrease in bottom-up learning and an increase in top-down learning, driven by the continuous maturation of the arcuate fasciculus, most likely provides the neurobiological basis for the gradual decrease of associative learning of NADs during early childhood, as observed in the current study.

Overall, our results are in line with previous studies arguing for different developmental stages of NAD learning (Culbertson et al., 2016) and add a new aspect: the notion of a gradual transition between these stages, supported by a reduction in amplitude and duration of the reported ERP effect. This gradual transition may point towards a sensitive period for associative NAD learning during early childhood. Under this view, younger children would have an advantage of learning NADs under passive listening conditions (Friederici et al., 2011; Mueller et al., 2019; van der Kant et al., 2020) compared to older children and adults (Friederici et al., 2013; Mueller et al., 2012; Mueller et al., 2019). This advantage would decrease gradually with age, as indicated by the linear decrease in our study. Older children and adults may still be able to learn NADs under passive listening in the presence of facilitating factors, such as additional acoustic cues (Frost & Monaghan, 2016; Gómez, 2002).

### 4.1 Limitations

As discussed in the introduction, previous studies have found different polarities of the ERP effect evoked in the present and similar NAD learning paradigms, that were related to different external variables, such as age (Friederici et al., 2011; Mueller et al., 2009), sex (Mueller et al., 2012), and children’s language development (Kooijman et al., 2013; Schaadt et al., 2015). Like these studies, we here found different ERP polarities of the NAD-related effect; however, none of the external variables we assessed (i.e., age, sex, language comprehension and language production) significantly explained this polarity difference. It is therefore not certain whether the two ERP responses (with positive and negative polarity) can be equated and the present results should be interpreted with caution in this regard. However, independent of whether the positive and negative ERP response indicate the same underlying process, we still consider the fact that both ERP amplitudes decrease with age informative, interpreted as an age-related decrease of the strength of NAD learning.

Moreover, our study is the first to find a *gradual* decrease of NAD learning under passive listening with age. While some form of age-dependent decrease of NAD learning under passive listening is well documented in the literature (Friederici et al., 2011; Mueller et al., 2009, 2019; van der Kant et al., 2020), more studies will be needed to confirm the gradual nature of this decrease. Here, we sampled NAD learning at three time points during early childhood, 1 year (13 months), 2 years (25 months), and 3 years (37 months) of age. Future studies will need to sample age more continuously to confirm the gradual decrease of NAD learning under passive listening conditions.

## 5 Conclusion

Our findings suggest a gradual decrease of associative NAD learning under passive listening during early childhood. Children at 1 to 3 years of age showed neurophysiological evidence of associative NAD learning under passive listening conditions, but the amplitude of this ERP effect linearly decreased with age. We propose that this linear decrease may be driven by entrenchment of children’s knowledge of their native language NADs, which may hinder NAD learning in a foreign language. In addition, brain maturation during early childhood likely contributes to children’s increasing ability to utilize higher-level, morphological features of the input through top-down learning, and to their decreasing ability to learn NADs associatively under passive listening conditions. Our study provides first evidence that the transition between different developmental stages of NAD learning may occur in a gradual manner, pointing toward a sensitive period for NAD learning during early childhood.

## Acknowledgements

The Max Planck Society (MP, CM, ADF), the German Research Foundation (DFG) in the Research Unit FOR 2253 (Project number 258522519), project FR 519/20-1 (MP, CM, AK, BH, IW, ADF), and the Berlin School of Mind and Brain (MP) funded this project. Further, we would like to thank all participating families for their commitment, as well as Anika Schütze, Bettina Nowotny, Bianca Dietrich, Christina Rügen, Claudia Geißler, Eileen Stelter, Kristiane Klein, Leonie Faulhaber, and Sophia Richter for their help with data acquisition and their dedicated work with the participating children and families.

