## Supplementary Materials for "Gradual development of non-adjacent dependency learning during early childhood"

### **Analysis including outliers**

We excluded outliers from our dataset, following two approaches. First, we applied exclusion criteria based on the quality of children’s EEG data (see main text, section 2.1) and based on the absolute ERP amplitude, for which outliers were defined as 2.5 times the median absolute cutoff (see main text, section 2.5.2). This procedure resulted in the exclusion of the datasets of 6 additional children. Here, we report the results of the linear mixed model before outlier exclusion. These results were similar to the linear mixed model after outlier exclusion, that is, all effects were on the same side of the significance threshold (p<0.05). In particular, the LM with outliers showed a significant effect of age (β = -2.07, p = 0.003; table S1). The follow-up LMs (equivalent to a one-sample t-tests including weights for the number of trials that constituted each average) for each age group revealed that, for all age groups, the ERP absolute ERP amplitude was significantly different from 0 (1 year: β = 6.42; p < 0.001; 2 years: β = 4.32; p < 0.001; 3 years: β = 3.48).

Table S1. Summary of the linear mixed model of children’s ERP amplitudes including outliers

| *Predictors* | *Estimates* | *CI* | *p* |
| --- | --- | --- | --- |
| (Intercept) | 4.73 | 3.95 – 5.51 | **<0.001** |
| age | -2.07 | -3.43 – -0.70 | **0.003** |
| Observations | 115 | | |
| R^2^ / R^2^ adjusted | 0.074 / 0.066 | | |

**Latency analysis**

Table S2. Full table of the results of the follow-up t-tests on the latency of the ERP effect for each age group. P-values (pvalues_corr) were Bonferroni-corrected. Significant p-values are displayed in bold font.

|  | 12-mo | | 24-mo | | 36-mo | |
| --- | --- | --- | --- | --- | --- | --- |
| TW | pvalues_corr | t-values | pvalues_corr | t-values | pvalues_corr | t-values |
| 0-100 | 0.415462099 | 2.25714278 | 11.5190394 | 0.22547323 | 8.78346043 | 0.48984426 |
| 100-200 | 0.096927608 | 2.85147127 | 0.52797723 | 2.15121386 | 4.30313885 | 1.03632111 |
| 200-300 | **0.006786692** | 3.80744477 | 2.75533704 | 1.31314011 | 0.1995691 | 2.58343966 |
| 300-400 | **0.000976224** | 4.45014352 | 7.10113973 | 0.66933647 | 0.09820444 | 2.86981968 |
| 400-500 | **8.98924E-06** | 5.93279956 | 0.12489933 | 2.75288089 | 0.05959546 | 3.0638681 |
| 500-600 | **4.98525E-06** | 6.11711461 | **0.00144405** | 4.32267004 | 0.11996529 | 2.79039979 |
| 600-700 | **4.2748E-07** | 6.88811719 | **1.9465E-05** | 5.69120888 | **0.0047398** | 3.98446818 |
| 700-800 | **1.09123E-06** | 6.59304843 | **0.00334204** | 4.04584517 | **0.000193** | 5.07368389 |
| 800-900 | **3.47828E-06** | 6.2297063 | **0.00067773** | 4.56815689 | **0.00019114** | 5.07693106 |
| 900-1000 | **0.000762352** | 4.53018788 | **0.00095585** | 4.45698491 | **0.00028085** | 4.9479835 |
| 1000-1100 | **0.010751435** | 3.64977139 | **0.06759137** | 2.98881329 | 0.51867163 | 2.17047061 |
| 1100-1200 | 0.512261213 | 2.16472839 | 0.43825973 | 2.233774 | 5.39744968 | 0.87907366 |
| 1200-1300 | 0.364661166 | 2.31364385 | 0.6995757 | 2.02297211 | 1.78442298 | 1.56238318 |
| 1300-1400 | 0.42438982 | 2.24785824 | 0.69391544 | 2.02673752 | 3.53795693 | 1.16354015 |
